# Driving Integrative Structural Modeling with Serial Capture Affinity Purification

**DOI:** 10.1101/2020.04.08.032151

**Authors:** Xingyu Liu, Ying Zhang, Zhihui Wen, Yan Hao, Charles A.S. Banks, Jeffrey J. Lange, Brian D. Slaughter, Jay R. Unruh, Laurence Florens, Susan M. Abmayr, Jerry L. Workman, Michael P. Washburn

## Abstract

Streamlined characterization of protein complexes remains a challenge for the study of protein interaction networks. Here, we describe Serial Capture Affinity Purification (SCAP) where two separate proteins are tagged with either the HaloTag or the SNAP-tag, permitting a multi-step affinity enrichment of specific protein complexes. The multifunctional capabilities of these protein tagging systems also permit *in vivo* validation of interactions using FRET and FCCS quantitative imaging. When coupling SCAP to cross-linking mass spectrometry, an integrated structural model of the complex of interest can be generated. We demonstrate this approach using the Spindlin1 and SPINDOC chromatin associated protein complex, culminating in a structural model with two SPINDOC docked on one SPIN1 molecule. In this model, SPINDOC interacts with the SPIN1 interface previously shown to bind a lysine and arginine methylated sequence of histone H3 Taken together, we present an integrated affinity purification, live cell imaging, and cross linking mass spectrometry approach for the building of integrative structural models of protein complexes.

## Introduction

Estimates of the number of human protein-protein interactions (HPPIs) continue to grow. In 2017, 625,641 HPPIs were predicted ^1^ while a subsequent estimate yielded nearly 1 million HPPIs ^2^ This number will likely grow considering that there are a vast number of cells in a human body and a myriad of different cellular conditions in normal and diseased states. New methods are needed to tackle the enormous challenge of validating and determining the structural and functional significance of proposed HPPIs and how they are organized into larger networks inside of cells.

One way to tackle this challenge is to first develop computational methods to identify potential direct protein-protein interactions ^3^, and then to use these predictions to guide further experimental studies. An established and important further experimental approach is affinity purification followed by mass spectrometry (APMS) in which an affinity-tagged protein is purified along with its interactors, which are then identified by liquid chromatography coupled to mass spectrometry (LCMS). This APMS approach has been successful but there are computational challenges associated with distinguishing non-specific interactions ^4^. Foundational work by Rigaut et al. described an AP approach for the study of *S. cerevisiae* protein complexes, where a protein was fused to two affinity tags enabling a two-step enrichment method resulting in protein complexes of higher purity ^5^. This TAP-tag method played a key role in the analysis of *S. cerevisiae* protein complexes and protein interaction networks ^6^. Recently, more advanced multifunctional tags such as the HaloTag ^7^ and SNAP-tag ^8^ have been developed, which can be used for both affinity purification and microscopy imaging methods.

Combining these concepts, we have devised a strategy to study any pair of proteins that might directly associate to 1) validate their interactions via proteomics, 2) assess their interactions in live cells, and 3) build a structural model of the complex. Here, we describe the development of reagents for Serial Capture Affinity Purification (SCAP), an approach fundamentally different from conventional affinity purification. SCAP uses a combination of two separately tagged bait proteins to reduce the complexity and increase the purity of protein complexes. SCAP can be followed by label-free quantitative mass spectrometry analysis (SCAP-MS) to identify protein complexes containing the two interacting proteins of interest. Furthermore, the SCAP constructs can be used to validate their interaction *in vivo* using quantitative imaging techniques. Finally, the SCAP pipeline can also include a cross-linking (XL) step followed by MS to identify cross-linked peptides defining interaction interfaces and intermolecular distance constraints, which can be used to build integrated molecular models of protein complexes.

We demonstrate the striking capabilities of this technology using a pair of proteins, Spindlin1 (SPIN1) and SPINDOC (c11orf84), which have previously been proposed to directly interact in biochemical ^9^ and computational ^3^ studies. SPIN1 is a well characterized histone methylation reader ^9–21^, while SPINDOC has only been defined by its ability to bind SPIN1 ^9^ In this study, we first characterize the direct interaction and co-diffusion of SPIN1 and SPINDOC in live cells using imaging methods. We next used SCAP-MS to generate a sample enriched with SPIN1 and SPINDOC. This fraction was then analyzed using advance XL techniques followed by molecular modeling. The culmination of these studies resulted in an integrated structural model of a complex formed by one molecule of SPIN1 and two molecules of SPINDOC.

## Results

### Building reagents for Serial Capture Affinity Purification

In a typical APMS study, a protein of interest (POI) is affinity tagged and transiently expressed in cells. The tagged POI is then used to capture protein complexes from cell extracts and proteins co-purifying with the POI are identified by mass spectrometry. Although this is a well-proven approach, it essentially analyses *ex vivo* complexes that might not reflect genuine interactions within a cell. As an example, we expressed Halo-SPIN1 (Figure S1A) and Halo-SPINDOC (Figure S1B) separately in HEK293 cells, affinity purified the associated proteins, and analyzed them by label free quantitative proteomics (Supplemental Table S1). The resulting data demonstrates that the bait protein was the most abundant protein in each sample with ~7X less SPINDOC than SPIN1 in the Halo-SPIN1 purification (Figure S1A) and ~20X less SPIN1 than SPINDOC in the Halo-SPINDOC purification (Figure S1B). There were many additional proteins in both purifications (Supplemental Table S1). There are several challenges in interpreting such APMS datasets: first, defining which proteins genuinely interact with the POI; second, determining which proteins interact directly with the POI; and third, establishing the stoichiometry of the proteins in purified complexes.

Considering such challenges, new approaches are needed to better characterize protein complexes in a streamlined and efficient manner. We therefore devised the SCAP approach using two orthogonal affinity tags: the HaloTag ^22^ and the SNAP-tag ^8^. These tags are multifunctional and facilitate multiple different types of analyses using a single construct. For example, these tags can be labeled with fluorophores for live cell imaging in addition to being used for affinity purification. Both the HaloTag and SNAP-tag covalently bind to their respective substrates immobilized on beads. There is an established system for elution of immobilized Halo tagged proteins isolated from mammalian cell extracts. A linker sequence between the Halo tag and the POI contains a TEV protease cleavage site^22–24^, allowing TEV protease mediated release of the POI and associated proteins from the beads, leaving the HaloTag bound to the beads ^22^ In contrast, a system for purifying SNAP-tagged proteins from mammalian cells is not well established.

To develop a SNAP purification strategy that could be used together with the Halo purification system, allowing independent cleavage of Halo and SNAP-tagged baits, we needed to choose a protease that recognized a cleavage sequence different from TEV protease. To evaluate the suitability of proteases to use for eluting SNAP isolated proteins, we constructed N’-terminal SNAP-tag versions of SPIN1 with several different protease cleavage sites (Figure 1A). We expressed these versions of SNAP-SPIN1 in HEK293/FRT cells, isolated SPIN1 using the SNAP-tag, and used the indicated protease to elute SPIN1 for analysis by Western blotting (Figure 1B). The quantity and purity of SNAP-SPIN1 isolated using PreScission protease were comparable to those of SNAP-SPIN1 isolated using TEV protease (Figure 1B). In contrast, enterokinase and Factor Xa protease have shorter recognition sequences than TEV and PreScission protease, and not only cleave the linker, but also appear to cleave SPIN1 (Figure 1B), limiting the utility of these proteases. Therefore, we chose a PreScission protease-based cleavage system as our elution method for SNAP-tag purification.

**Figure 1.**
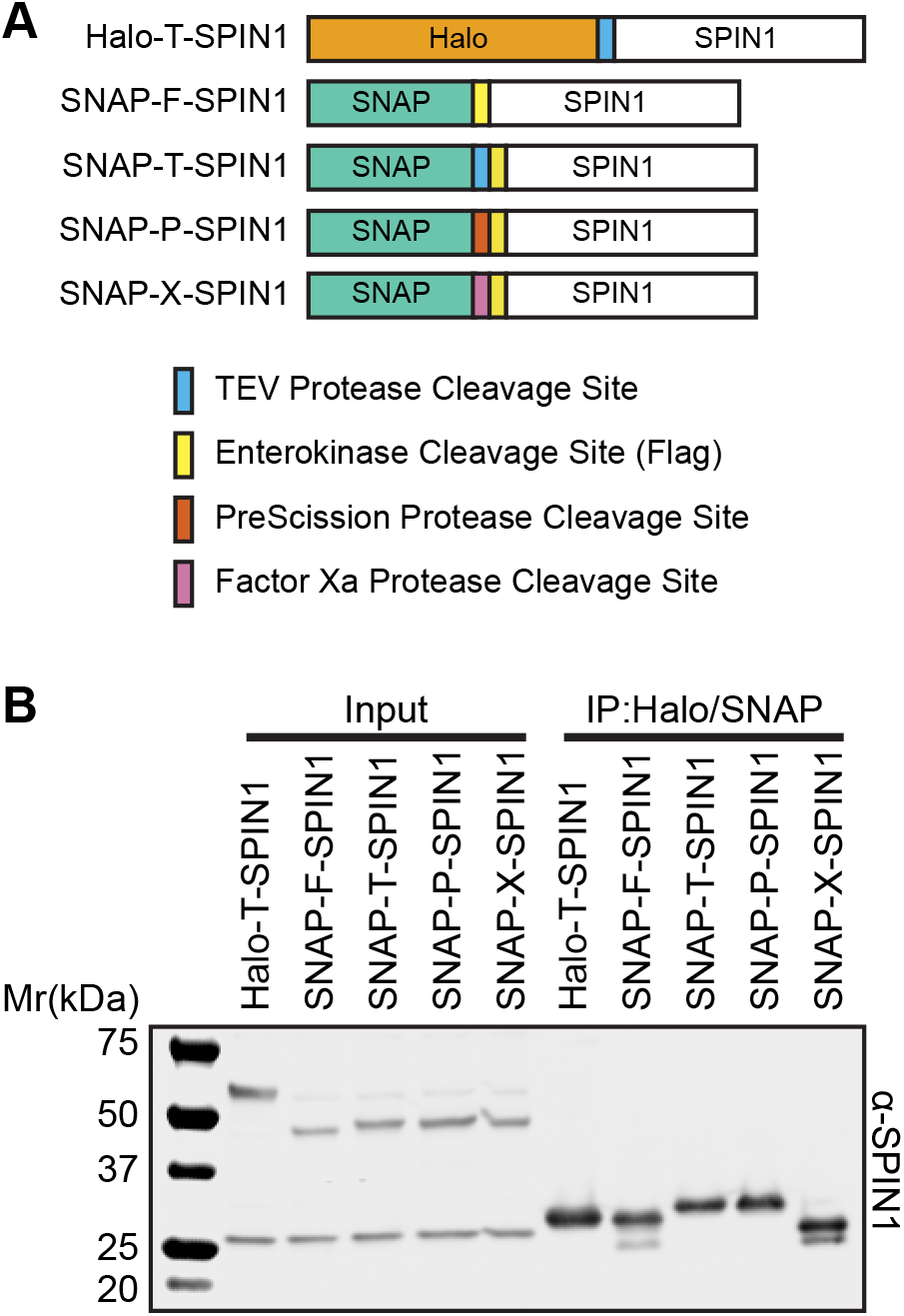
Development of Constructs for SNAP-AP. (A) Schematic of constructs of HaloTag and SNAP-tag SPIN1 with different protease recognition sites. (B) Anti-SPIN1 western blot where HaloTag and SNAP-tag SPIN1 proteins were transiently transfected into HEK293FRT cells. Whole cell lysates were collected as the input sample and Halo or SNAP purifications were performed followed by SPIN1 elution from beads using the corresponding protease as the IP sample.

### Developing a Serial Capture Affinity Purification

To perform SCAP, we first constructed an expression vector based on pcDNA5/FRT that would enable us to express both a Halo- and a SNAP-tagged protein from the same plasmid. We inserted sequences coding for both the HaloTag and SNAP-tag each followed by convenient restriction sites for subcloning our two bait proteins. (Figure 2A and Figure S2A). As our plasmid generates an mRNA coding for two tagged proteins, we also engineered an internal ribosomal entry site (IRES) between the sequences coding for each protein. We then inserted the open reading frames for the two proteins of interest into the ORF1 and ORF2 regions. These plasmids can be used for either transient or stable expression. To develop a sequential purification system, we next generated a stable cell line that expressed Halo-SPIN1 and SNAPSPINDOC (Figure 2B). First, proteins from whole cell extracts prepared from these cells were isolated on SNAP affinity beads and then eluted using PreScission protease (fraction E1). We next used 80% of fraction E1 for further affinity purification of tagged complexes using Halo affinity beads, retaining 20% for mass spectrometry analysis. The unbound supernatant of the Halo purification was collected as fraction UB2. The proteins captured by the Halo affinity beads were eluted using the TEV protease as fraction E2. Proteins from the three fractions E1, UB2, and E2 were analyzed by silver-stained SDS-PAGE (Figure 2C). This analysis clearly indicated a more complex protein mixture in the E1 and UB2 fractions compared with fraction E2. The analysis of fraction E2 generated two major bands consistent with enrichment of SPIN1 and SPINDOC proteins, a minor band (25 kDa) consistent with TEV protease, and two other minor unidentified bands (~70 kDa). The SCAP purification thus generated a sample with a high concentration of the interacting proteins of interest, SPIN1 and SPINDOC, removing most contaminants.

**Figure 2.**
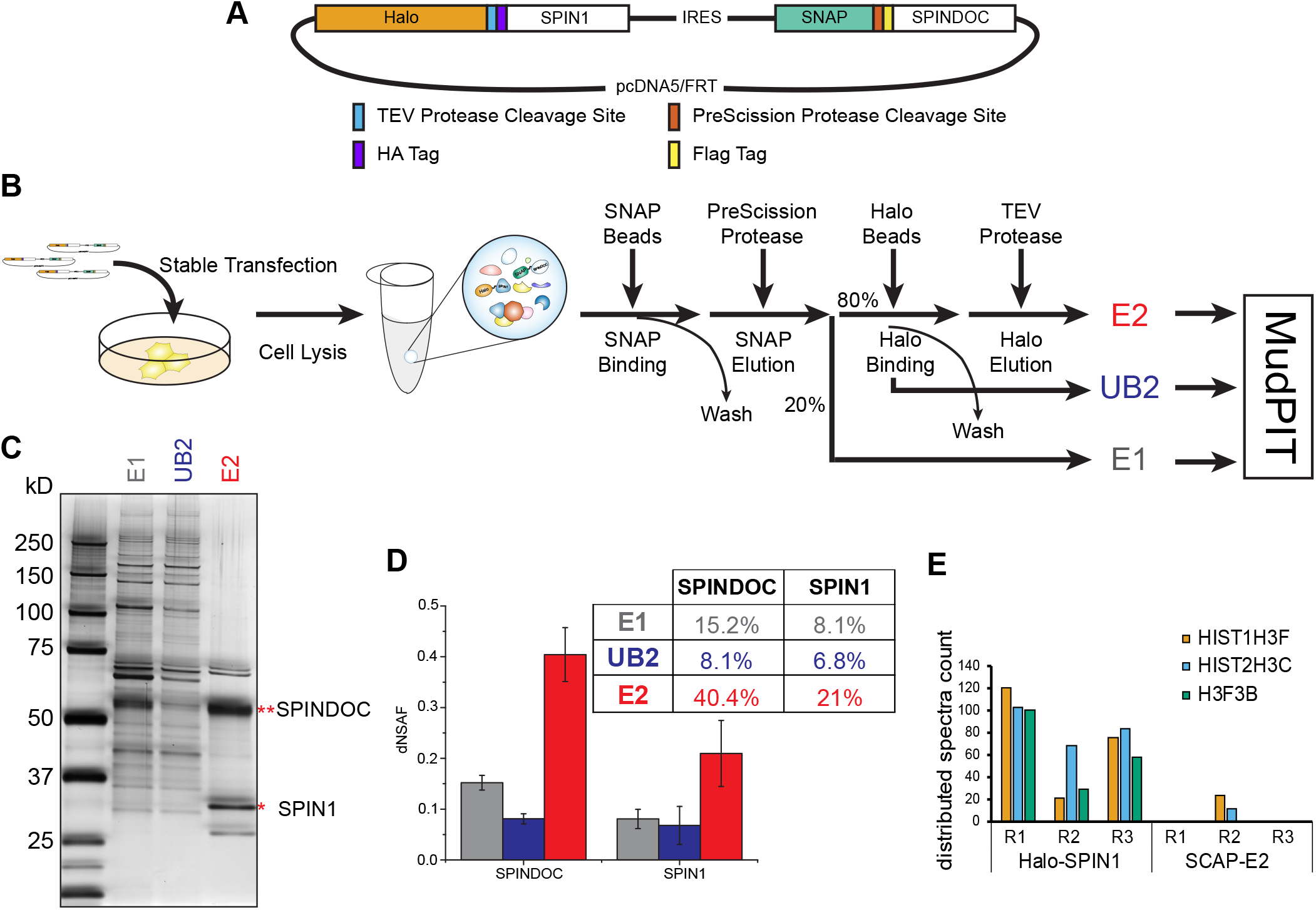
Serial Capture Affinity Purification of SPIN1 and SPINDOC Complexes. (A) Schema of the expression vector designed for co-expression of HaloTag and SNAP-tag protein pairs, with the detailed vector map provided in Figure S2A. (B) Workflow of Serial Capture Affinity Purification quantitative proteomics method. E1 (the elution from SNAP purification, UB2 (the unbound proteins after binding to Halo bead resin), and E2 (the elution from the Halo beads) were all separately analyzed by MudPIT and dNSAF values calculated for all identified proteins. The abundance of SPIN1 and SPINDOC were calculated as dNSAF x 100%. (C) Silver stained SDS-PAGE of the proteins eluted from the E1, UB2, and E2 fractions. (D) The dNSAF plot of SPINDOC and SPIN1 in E1, UB2, and E2 fractions. (E) Spectral counts measured for histones associated with SPIN1 in a single Halo purification and after SCAP.

To confirm the presence of SPIN1 and SPINDOC and assess their enrichment at each stage of the purification, all three E1, UB2, and E2 fractions were subjected to label free quantitative proteomic analysis using MudPIT (Supplemental Table S2). The averaged dNSAF values of the top 20 proteins in each fraction are shown in Figure S2B-D. An overall comparison of the enrichment of the SPINDOC and SPIN1 pair in each fraction is summarized in Figure 2D. Over the course of the 2-step SCAP protocol, the enrichment of SPINDOC and SPIN1 increased more than 2.5 fold: after SNAP purification, the spectral counts matching these two proteins contributed 15.2% and 8.1% of the total spectral counts, respectively, while after the 2^nd^ purification step, their enrichment was measured as 40.4% and 21%. In both E1 and E2 eluates (Figure 2D), the ratio dNSAF_SPINDOC_:dNSAF_SPIN1_ was approximately 2:1, suggesting a stoichiometry of two SPINDOC molecules for each SPIN1 molecule.

### Assessing the interaction of SPIN1 and SPINDOC in live cells

Next, using the multifunctional capability of the Halo and SNAP tagging systems, we analyzed the interaction of Halo-SPIN1 and SNAP-SPINDOC in live cells (Figure 3). To confirm the interaction between SPIN1 and SPINDOC in vivo, we implemented two different imaging-based approaches: Acceptor Photobleaching Förster Resonance Energy Transfer (AP-FRET) ^25^ and Fluorescence Cross-Correlation Spectroscopy (FCCS) ^26^. Both techniques can be applied in live cells, and both benefit from low concentrations of labeled proteins expressed in cells. These features reduce the possibility that SPIN1 capture of SPINDOC is due either to overexpression of the bait protein or to breakdown of subcellular segregation during cell lysis.

**Figure 3.**
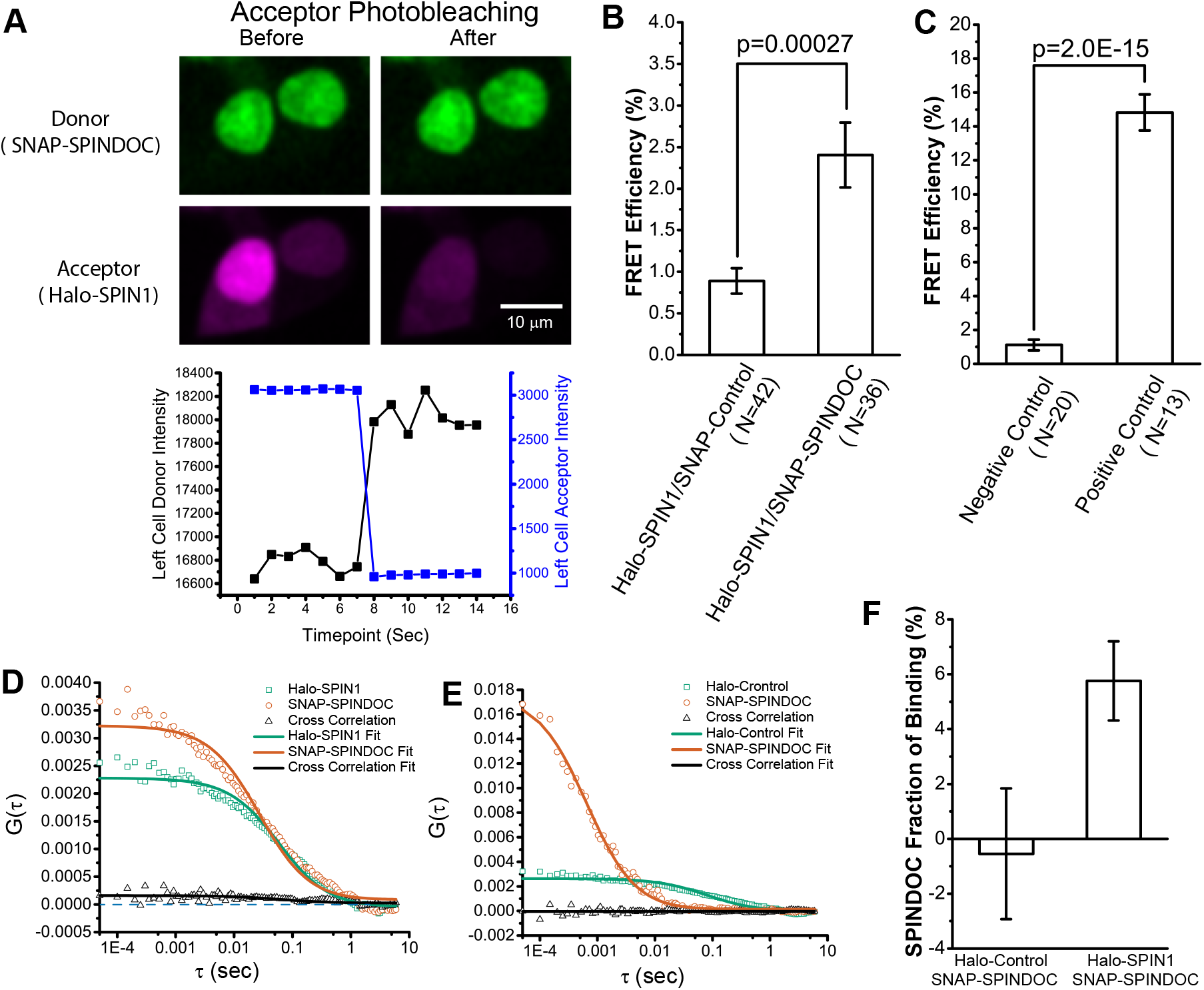
SPIN1 and SPINDOC Interaction in Live Cells. (A) Example image and intensity measurement of acceptor-photobleaching (AP) fluorescence resonance energy transfer (FRET). HaloTag SPIN1 was labeled with HaloTag TMRDirect and SNAP-tag SPINDOC tag was stained with SNAP-Cell 505-Star ligand. (B) Averaged FRET efficiencies measured for Halo-SPIN1 and SNAP-SPINDOC in live HEK293FRT cells. (C) FRET efficiencies measured for control proteins in live HEK293FRT cells. HaloTag and SNAP-tag alone were used as a negative control pair and a fusion protein with both the HaloTag and SNAP-tag was used as positive control. For both (B) and (C) bar charts, error bars stand for standard error of means for the datapoints defined in Figure S3A-B and p-values were calculated using two-tailed t-test. Fluorescence Cross-Correlation Spectroscopy of auto and cross-correlation curves for Halo-SPIN1 and SNAPSPINDOC (D) and for Halo-Control with SNAP-SPINDOC (E). (F) The average percentage of SNAP-SPINDOC binding to HaloTag control or Halo-SPIN1 was calculated from the y amplitudes of correlation curves. Error bars stand for standard error of the mean.

To apply AP-FRET on SPIN1 and SPINDOC, we co-expressed a Halo-SPIN1 and SNAP-SPINDOC in HEK293/FRT cells. Then we labeled the HaloTag with TMRDirect ligand as an acceptor and the SNAP-tag with 505Star ligand as the donor. We photobleached the acceptor cell by cell with a 561nm laser and measured the average donor intensity change of each bleached cell before and after bleaching of the acceptor (Figure 3A). The FRET efficiency of each cell was then calculated from the increased donor intensity after photobleaching of acceptor to the donor intensity after bleaching; multiple cells were analyzed (Figure S3B and Supplemental Table S3), and the average FRET efficiency was calculated (Figure 3B).

Similarly, protein pairs serving as positive and negative controls were also tested to determine the upper limit of FRET efficiency (Figure 3C and S3A). Halo-SPIN1 showed a significantly higher FRET efficiency with SNAP-SPINDOC than with SNAP itself (Figure 3B), indicating a direct interaction between SPIN1 and SPINDOC in live cells.

Next, we used another tool for detecting direct interactions, FCCS ^26^, to investigate whether Halo-SPIN1 and SNAP-SPINDOC co-diffuse with each other in live cells (Figure S3C). As with AP-FRET, we co-expressed SNAP-SPIN1 with Halo-WDR76 or Halo-tag only in live cells. Then we labeled the two tags with ligands conjugated to distinct fluorophores and measured both the auto self-correlation function of each species and the cross-correlation functions between HaloTag and SNAP-tag (Figure 3D). From the curves, we observed a crosscorrelation of Halo-SPIN1 and SNAP-SPINDOC but not between Halo-Control and SNAPSPINDOC (Figure 3E). The fraction of Halo-SPIN1 binding to SNAP-SPINDOC was then calculated using G(τ)the respective amplitudes of the auto and cross-correlation functions (Figure 3F) and showed a significantly larger fraction of SPINDOC binding to SPIN1 than to the HaloTag by itself. These results suggest that Halo-SPIN1 and SNAP-SPINDOC co-diffuse, and therefore interact in live cells.

### Implementing SCAP-XL to derive a structural model of the SPINl:SPINDOC complex

With the significant enrichment of the SPINDOC and SPIN1 proteins in the E2 fraction (Figure 2D), we reasoned that this highly purified complex would be an excellent candidate for further structural analysis. Taking advantage of the availability of MS-cleavable cross-linking reagents and highly sensitive mass spectrometers ^27,28^, we added a chemical cross-linking (XL) step to further improve the SCAP pipeline (SCAP-XL). The MS-cleavable disuccinimidyl sulfoxide (DSSO) cross-linker ^27^ was added to the SCAP isolated proteins bound to Halo beads before TEV protease elution, resulting in a cross-linked E2 fraction (Figure 4A). Analysis of the proteins on SDS-PAGE confirmed the presence of higher molecular weight cross-linked species (Figure S4A). We then analyzed these fractions in quadruplicate on an Orbitrap Fusion Lumos Tribrid mass spectrometer where cross-linked peptides were identified using MS1, MS2, and MS3 information (Figure S4B). The resulting mass spectrometry datasets were analyzed with the XlinkX search engine implemented through Proteome Discoverer ^29^ (Supplemental Table S4). The locations of the intermolecular cross-links between SPINDOC and SPIN1 and the locations of intramolecular cross-links were plotted using xiNET ^30^(Figure 4B). The structure of SPIN1 had been previously solved by X-ray crystallography ^31–33^. However, the structure of its dominant interacting protein, SPINDOC, remains to be elucidated, and the structural nature of the SPINDOC:SPIN1 complex is not well understood.

**Figure 4.**
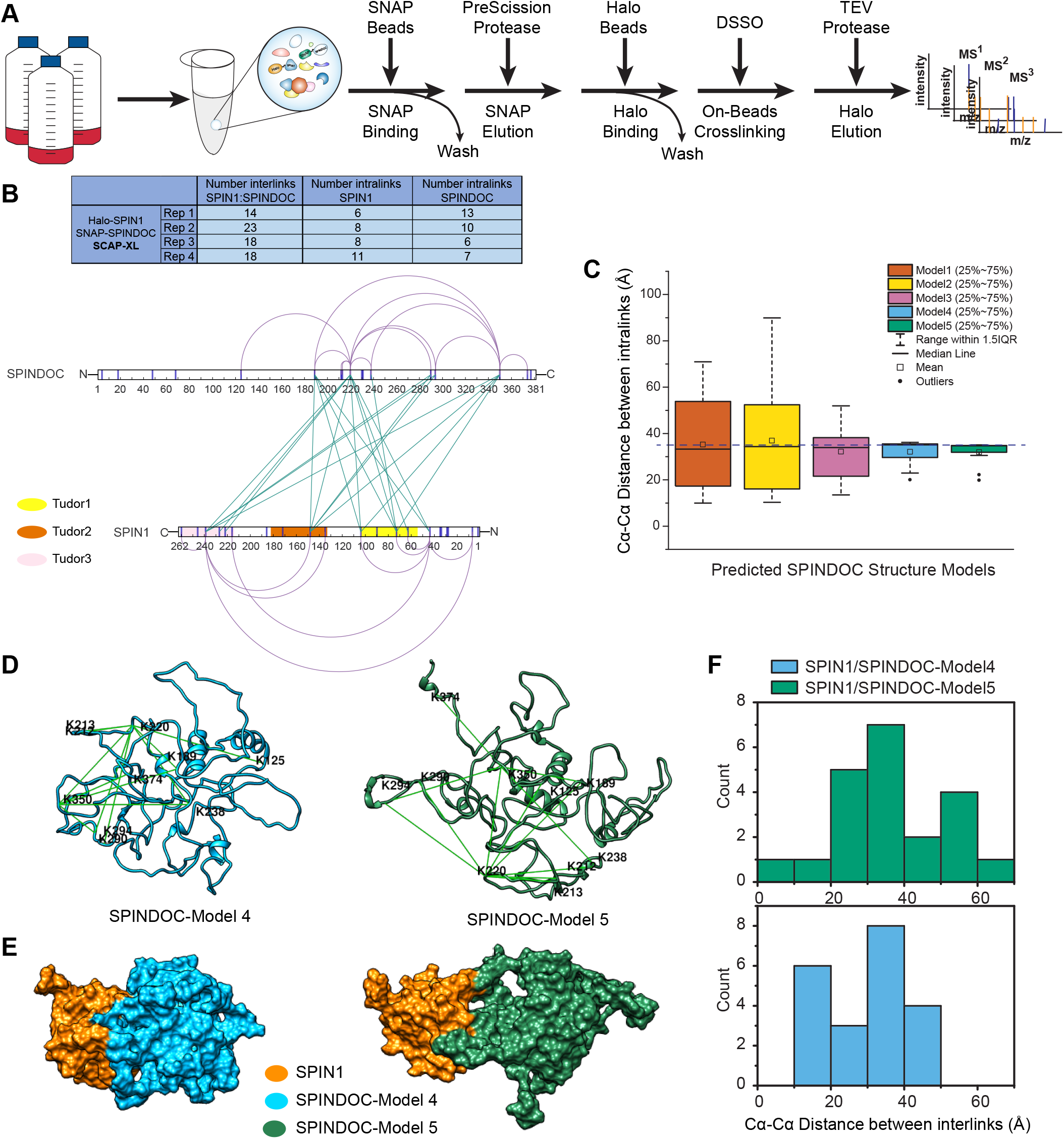
Integrated Modeling of Complex with Serial Capture Affinity Purification Coupled with Cross-linking. (A) Workflow of SCAP coupled with Cross-linking (SCAP-XL) method where a disuccinimidyl sulfoxide (DSSO) cross-linking reaction was performed before the TEV protease elution while the purified proteins were still on Halo beads. (B) A twodimensional visualization of SPIN1 and SPINDOC cross-links via XiNET ^30^. Self-cross-links are shown in purple and intermolecular cross-links are shown in blue. The total number of crosslinks detected and identified in replicate SCAP-XL analyses are shown in the table inset (See Supplemental Table S4 for details). (C) Cα-Cα distances between SPINDOC self-cross-link sites in structural models predicted by I-Tasser ^34^ (D) SPINDOC structural models 4 and 5 defined by I-Tasser ^34^ Self-cross-links are shown as green lines. (E) Docking models of SPIN1/SPINDOC-model 4 and SPIN1/SPINDOC-model 5 generated by HADDOCK ^37^ (F) Cα-Cα distances between SPIN1/SPINDOC inter-cross-linked sites for models 4 and 5.

To gain a better understanding of SPINDOC:SPIN1 complex architecture, we first used I-Tasser ^34,35^ to generate structural predictions based on the SPINDOC amino acid sequence, with or without the intramolecular SPINDOC cross-links as distance constraints. Two sets of five structural models were generated through this computational process. We next measured the distances between the Cα of linked pairs of lysine residues in each of the predicted SPINDOC structural models and analyzed the distribution of these distances (Figure 4C and S4C-D). Compared to models generated without any cross-linking input (Figure S4D), of the five models generated two models were obtained where distance constraints from cross-linking were further considered in the modeling computation (Figure 4C/S4C). SPINDOC models 4 and 5 had the smallest ranges between interlinks in agreement with the predicted estimated distance allowed by the DSSO spacer arm (35Å) and were therefore selected for further analysis (Figure 4D).

Next, a SPIN1 structure (4MZF) ^33^, SPINDOC structural models, and cross-linked sites between the two proteins were subjected to analysis using HADDOCK, a web based complex modeling server ^36,37^ The 4MZF structure ^33^ was chosen since it is a structure of SPIN1 bound to H3-K4me3-R8me2 peptide, which therefore represents SPIN1 interacting with a portion of an additional protein. For SPINDOC, Model4 and Model5 were submitted separately resulting in two distinct models of a binary complex (Figure 4E). Next, the distances between each pair of inter-linked sites between SPINDOC and SPIN1 were measured in each docking model. The docking model of SPIN1 and SPINDOC Model4 had a distribution of distance between interlinks that better matched the DSSO predicted limits (Figure 4F).

The interface between SPIN1 and SPINDOC in this model and the cross-links between the two proteins are shown in more details in Figure 5A, with a 180° rotation shown in Figure 5B. In this model of the SPIN1:SPINDOC heterodimer, SPIN1 had two groups of intermolecular cross-links that mapped to distinct regions of SPINDOC. One group contained interlinks under 35Å (solid lines, Figure 5B), while in the other group (dashed lines), the interlinks were above 35Å (Figure 5A-B). This suggested that a binary model was not the optimal solution supported by the interlink data. Consistent with this, a stoichiometry of two SPINDOC molecules to one SPIN1 molecule in the complex had previously been indicated by the SDS-PAGE and quantitative proteomics analysis of the E2 fraction in the SCAP purification (Figure 2C-D). We therefore re-ran the model with this input via HADDOCK ^36,37^ to refine the structural model of the complex allowing for two SPINDOC molecules to dock one SPIN1 molecule (Figure 5C). The distances measured between Cα-Cα of interlinked lysine residues in this updated model of a compact heterotrimer are in better agreement with DSSO distance constraints (Figure 5D).

**Figure 5.**
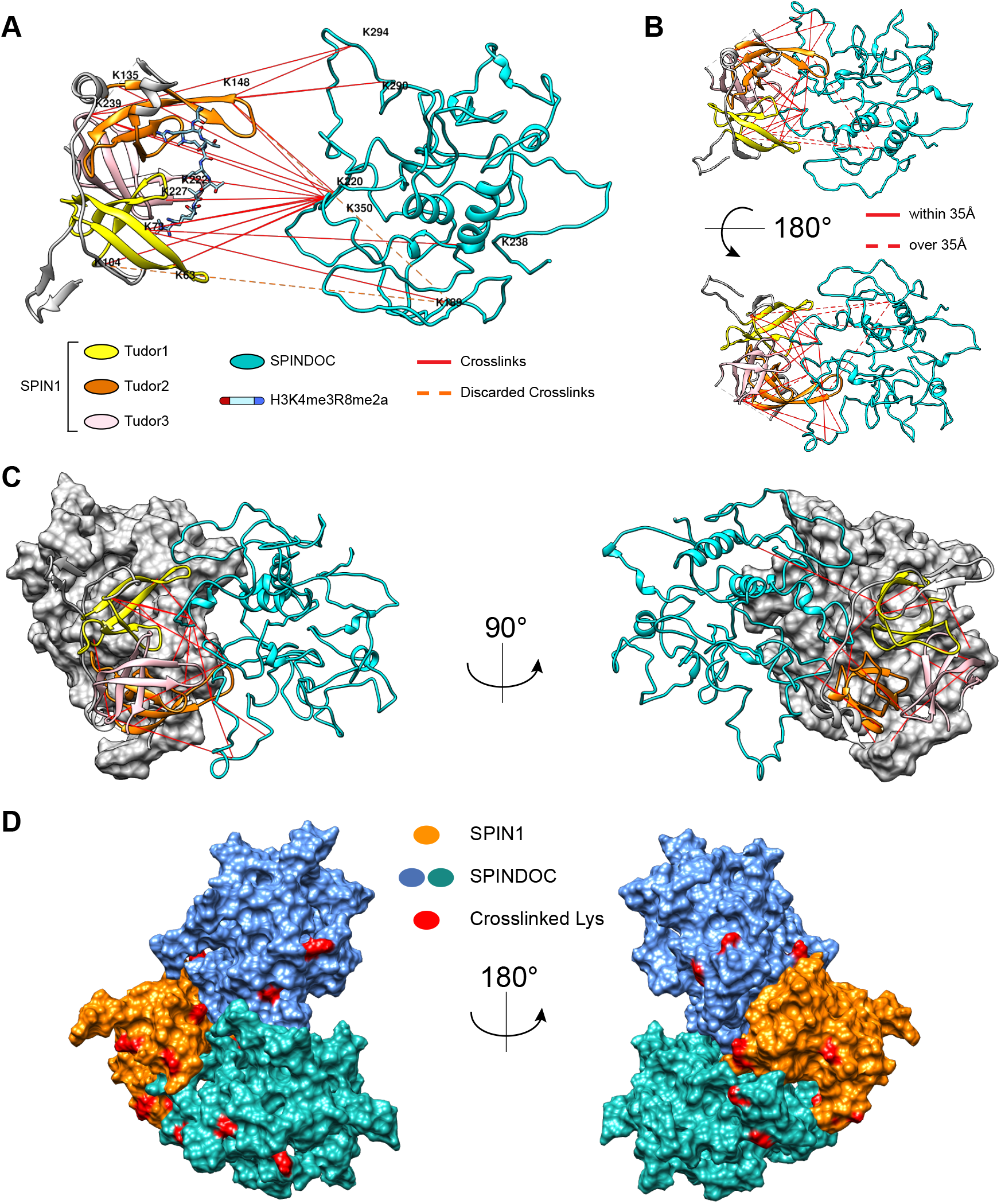
Integrative Modeling of the SPIN1: SPINDOC Complex. (A) Visualization of the intermolecular cross-links between SPIN1 and SPINDOC-model 4. (B) A HADDOCK-generated docking model with one copy of SPIN1 and one copy of SPINDOC-model 4. (C) Docking model with one copy of SPIN1 and two copies of SPINDOC-model 4. (D) Surface view of the quaternary structure of an heterotrimeric SPIN1:SPINDOC complex.

## Discussion

We designed SCAP by taking advantage of two multifunctional and orthogonal affinity tags, the HaloTag ^22^ and SNAP-tag ^8^ and, as a proof of principle, built a cell line stably expressing both Halo-SPIN1 and SNAP-SPINDOC for characterization in multiple experiments. The tudor domain containing protein SPIN1 is histone methylation reader and has been found to specifically bind H3K4me3 containing peptide with high affinity ^31–33,38^. In addition, SPIN1 has been reported to promote cancer proliferation and progression ^20,39^ The structure of SPIN1 has been solved by X-ray crystallography ^31–33^. As a result, multiple researchers are pursuing the development of a SPIN1 inhibitor ^10,16,40–43^. C11orf84, which was recently renamed SPINDOC by Bae and colleagues ^9^, is a SPIN1-interacting protein that is less well understood, and the complex containing these two proteins remains poorly characterized.

In a multi-step sequential affinity purification scheme, we named SCAP, proteins associated with SNAP-SPINDOC were first isolated on SNAP beads, then the population of such proteins also associating with Halo-SPIN1 were further enriched on Halo beads. An important feature of this approach is the use of distinct proteases for the Halo and SNAP purification steps where PreScission Protease was used to elute from the SNAP beads and TEV was used to cleanly elute proteins from the Halo beads. This protocol resulted in a more than 2.6-fold increase in both proteins concentration after elution from the 2^nd^ affinity step. The quantitative proteomic analysis of the final elution enriched in the complex suggested a 2:1 ratio between SPINDOC and SPIN1 molecules. The multifunctional features of the HaloTag ^22^ and the SNAPtag ^8^ were also used to validate the interaction between SPIN1 and SPINDOC with imaging approaches in live cells using the same expression constructs used for protein purification.

Upon optimization of the purification protocol for serial capture, the enriched population of Halo-SPIN1 and SNAP-SPINDOC complexes was the ideal candidate for further structural characterization using state of the art cross-linking mass spectrometry and computational approaches ^27,28^. We therefore used SCAP-XL to purify DSSO-cross-linked protein complexes for analysis on an Orbitrap Fusion Lumos mass spectrometer, which has advanced capabilities for the study of CID-cleavable cross-linked peptides ^28^. The culmination of this analysis was first defining a reliable tridimensional model for SPINDOC, for which no structural information is available, and second, refining a structural model of a heterotrimer, in which two SPINDOC molecules are docked on one SPIN1 molecule.

The SPIN1 structure we used to assemble the complex contained a histone H3-K4me3-R9me2a peptide ^33^, thus serving as a model for SPIN1 interacting with other proteins. Intriguingly, the SPINDOC:SPIN1 interaction surface identified by cross-linking overlaps with the binding pocket for H3-K4me3-R8me2a (Figure 5A). This result suggests that the binding of SPIN1 to SPINDOC could disrupt and/or compete with its binding to modified histone H3, which is consistent with previous findings of Bae and colleagues ^9^ Furthermore, when comparing the proteins recovered by proteomics analyses of the SCAP E2 elution to a Halo-SPIN1 purification alone, there were significantly reduced histone H3 interactions in the SCAP E2 elution (Figure 2E), also supporting the possibility that SPINDOC disrupts SPIN1 interaction with histone methylation sites, which warrants further study.

The SCAP and SCAP-XL pipelines described herein were designed to be generic approaches that can realistically be applied to any pair of interacting proteins. We devised a plasmid-based system for making stable cell lines in HEK293 cells. This enabled us to generate enough starting material for the SCAP-XL pipeline since cross-linked peptides can be of low abundance in a sample. This system could allow for medium throughput analysis of predicted direct protein interactions that are part of protein complexes of varying size ^3^, and these predicted directly interacting proteins are likely good candidates for incorporation into the SCAP and SCAP-XL pipelines for *ex vivo* complex characterization, *in vivo* interaction validation, and the building of structural models of protein complexes. The larger concept of ProteoCellomics is defined by coupling, on the one hand, quantitative proteomic analysis of affinity purified complexes and, on the other hand, quantitative spectroscopy techniques to image these complexes in live cells (Figure 6). SCAP is therefore a method enabling the practical application of ProteoCellomics, where quantitative proteomics, an *ex vivo* approach, and quantitative microscopy, an *in vivo* approach, are integrated to gain molecular insights into protein-protein interactions. Lastly, when then utilizing the SCAP-XL pipeline and computational modeling approaches an integrated structural model of a protein complex can be generated further advancing the understanding of poorly characterized protein protein interactions and protein complexes.

**Figure 6.**
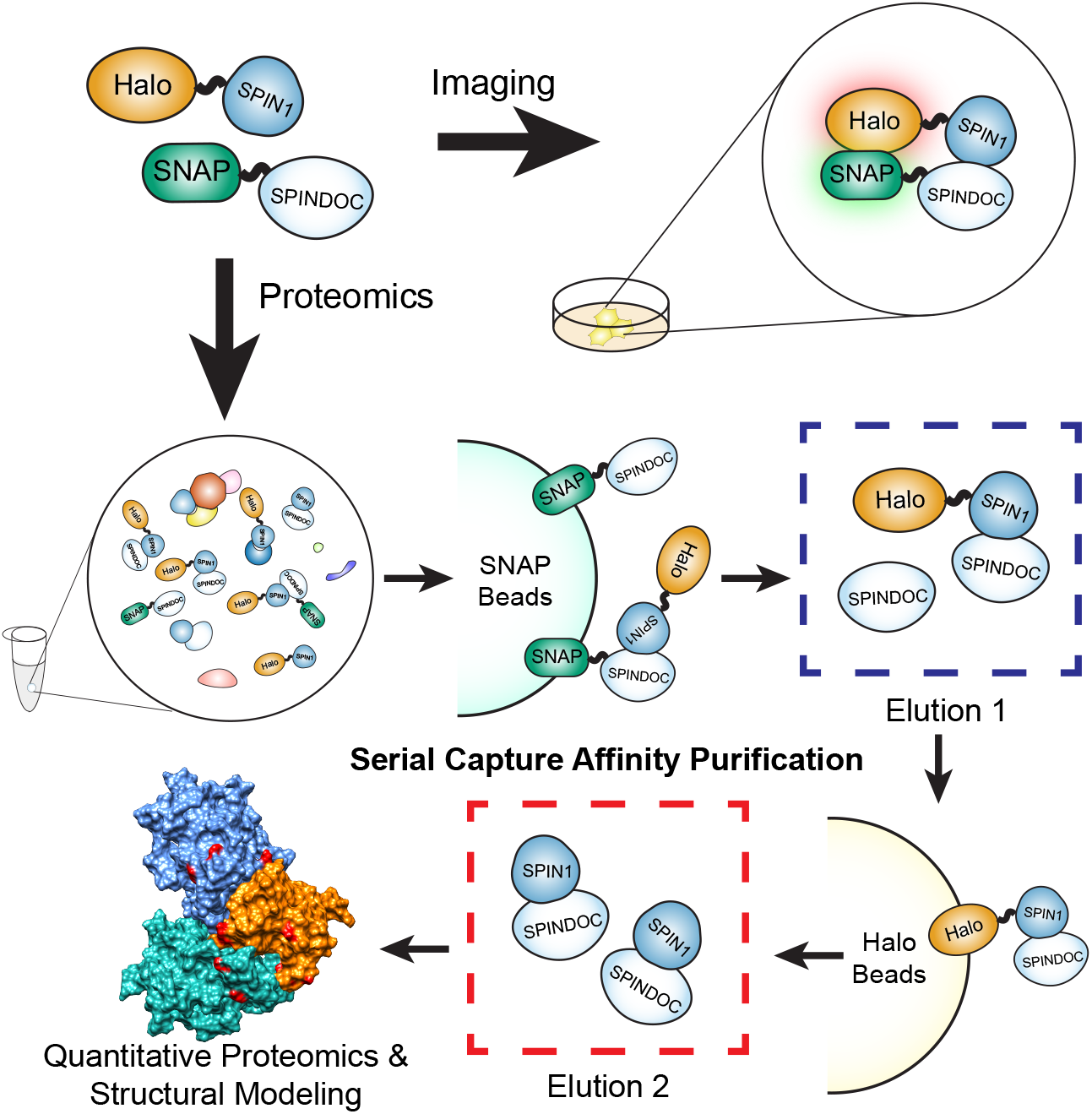
General concept of ProteoCellomics. In ProteoCellomics, protein interactions are characterized *ex vivo* using affinity purification and protein mass spectrometry, and *in vivo* using live cell imaging. This is accomplished by tagging two separate proteins with distinct multifunctional affinity tags like the HaloTag and the SNAP-tag. The interaction of two proteins can then be studied in a live cell using imaging techniques. Enriched protein complexes can then be isolated using Serial Capture Affinity Purification (SCAP) and analyzed using protein mass spectrometry techniques.

## Online Methods

### Critical Reagents

Magne^®^ HaloTag^®^ Beads, Sequencing Grade Modified Trypsin, rLys-C and HaloTag^®^ Ligands were purchased from Promega (Madison, WI, USA). All restriction endonucleases, SNAP-Cell^®^ ligands and SNAP-Capture Magnetic Beads were purchased from New England BioLabs (Ipswich, MA, USA). PreScission Protease was purchased from GE Healthcare Life Sciences (Chicago, Il, USA). AcTEV™ Protease and DSSO (disuccinimidyl sulfoxide) were purchased from Thermo Fisher Scientific (Waltham, MA, USA). Salt Active Nuclease (SAN) was purchased from ArcticZymes (Tromso, Norway).

### Plasmids and Cell lines

Sequences of SPIN1 open reading frame was obtained from Kazusa Genome Technology (Kisarazu, Chiba, Japan). Sequence of SPINDOC open reading frame was obtained from RT-PCR using mRNA extracted from 293FRT cells. Vectors containing SNAP-tag with different protease recognition sites were modified from pSNAPf (NEB). SPIN1 coding sequence were sub-cloned into pSNAPf-F/T/P/X vector for transient expression of SNAP-F/T/P/X-SPIN1. pcDNA5FRT vector was purchased from Invitrogen. pcDNA5FRT-Halo was generated by inserting Halo-TEV sequence (obtained from pFN21A vector purchased from Promega) downstream of CMV promoter. pcDNA5FRT-SNAP was generated by inserting SNAP-PP sequence (obtained from pSNAP-P vector) downstream of CMV promoter. Positive control vector pcDNA5FRT-Halo-NLS-SNAP was generated by linking Halo and SNAP tag sequence by an NLS sequence and inserted downstream of CMV promoter of pcDNA5FRT vector. The duel expression vector was modified from pcDNA5FRT-Halo vector by inserting internal ribosome entry site (IRES) sequence downstream of Halo-TEV-HA sequence, then followed by SNAP-PP-Flag sequence. Then SPIN1 or SPINDOC sequences were sub-cloned into each vector for expression of corresponding Halo or SNAP tagged protein. All oligos used in cloning were synthesized by Integrated DNA Technologies (Coralville, IA, USA).

293FRT cells were purchased from Invitrogen (Carlsbad, CA, USA) and maintained in DMEM medium with GlutaMAX and 10% FBS. All stable expression cell lines were maintained in DMEM medium with GlutaMAX, 10% FBS and 100 ug/mL hygromycin B. For transient expression, FuGene6 (Promega) was used to transfect expression vector into 293FRT cells. Stable expression cell lines were generated using Flp-In System according to manual (Invitrogen). All vectors we have mentioned above contain CMV promoter to drive expression. All stable expression cell lines used in this manuscript was generated from 293FRT cells. All cells were maintained at 37°C in 5% and humidified incubator.

### Single-Bait Purification

Halo-SPIN1 purification was performed using two different clones of Halo-SPIN1 stable expression cell lines as material. 293FRT cells were used as control cells. Both Halo-SPIN1 expression cells and control cells were subjected to Halo purification according to the manual of HaloTag^®^ Mammalian Pull-Down Systems (Promega). Eluates for a total of 6 replicates of SPIN1 purification (3 for each clone) and 3 replicates of control purification were subjected to MudPIT analysis. For control and Halo-SPINDOC purifications, pcDNA5FRT-Halo-NLS-SNAP or pcDNA5FRT-Halo-SPINDOC was transfected to 293FRT cells. Cells were collected 48hr after transfection and subjected to Halo or SNAP purification. SNAP purification protocol was similar to Halo purification, except for using SNAP-Capture Magnetic Beads (NEB) instead of Magne^®^ HaloTag^®^ Beads (Promega) and using PreScission Protease instead of TEV Protease to elute. Three replicates of Halo control purifications and 3 replicates of Halo-SPINDOC purifications were analyzed by MudPIT.

### SCAP and SCAP-XL

For regular SCAP, both baits were stably expressed in 293FRT cells. Cells were collected and lysed with High Salt Lysis Buffer. The lysate was centrifuged, and supernatant was incubated with SNAP-Capture Magnetic Beads (NEB) at 4°C for 2hr. Beads were then washed with High Salt Wash Buffer for 3 times followed by Wash Buffer for 2 times. Bound proteins were eluted with Elution buffer (containing PreScission Protease). 20% of the eluate was aliquoted as E1 and the rest 80% was subjected to further Halo purification step. Eluate obtained above was then incubated with Magne^®^ HaloTag^®^ Beads (Promega) at 4°C for 2hr. Unbound supernatant was collected as UB2. Beads were then washed with Wash Buffer for 5 times. Bound proteins were eluted with Elution buffer (containing TEV protease) and the eluate is collected as E2. For all three replicates, E1, UB2 and E2 are subjected to MudPIT analysis. For SCAP-XL, cells were collected from 3 roller bottles. Cell pellet was homogenized by dounce tissue grinder in high salt lysis buffer and lysed at 4°C. Lysate was centrifuged and supernatant was subjected to SNAP purification. Then the eluate was bound to Magne^®^ HaloTag^®^ Beads (Promega). After washes, bound proteins were cross-linked on beads with 5mM DSSO (Thermo Fisher) at 4°C for 1hr. The cross-linking reaction was quenched by 50mM Tris-HCl and crosslinked proteins were eluted by TEV protease.

### Mass Spectrometry Sample Preparation

5% of each sample was used for SDS-PAGE and silver staining analysis before processed for mass spectrometry analysis (data not shown). Each sample was first TCA precipitated and then resuspended with 8M Urea buffer (in 100mM Tris-HCl, pH 8.5). The resuspended proteins were reduced with tris(2-carboxyethyl)phosphine (TCEP) and treated with 2-Chloroacetamide (CAM). Then proteins were digested with Lys-C for at least 6 hours followed by overnight trypsin. Last, the digested samples were quenched with formic acid before subjected to MudPIT analysis ^44^.

### Multidimensional Protein Identification Technology (MudPIT)

MudPIT has been described before ^45^. Each digested sample was loaded onto a three-phase column. The column was pulled from capillary (100 μm i.d.) to a 5 μm tip, then packed first with 8 cm of 5 μm C18 RP particles (Aqua), followed by 3.5 cm of 5 μm Luna SCX, and last with 2.5 cm of 5 μm Aqua C18. Then the loaded column was washed with Buffer A (5% Acetonitrile, 0.1% Formic Acid) before placed onto instruments. For Halo-SPIN1 single-bait purification samples and control samples, each loaded column was placed in line with an Agilent 1100 quaternary HPLC pump (Palo Alto, CA) and an LTQ mass spectrometer (Thermo Fisher Scientific). Each full MS scan (400-1600 m/z) was followed by five data-dependent MS/MS, the number of microscans was 1 for MS and MS/MS scans. For SCAP samples, each loaded column was placed in line with an Agilent 1200 quaternary HPLC pump (Palo Alto, CA) and a Velos Orbitrap Elite mass spectrometer (Thermo Fisher Scientific). MS spray voltage set at 2.5 kV; MS transfer tube temperature set at 275°C; 50 ms MS1 injection time; 1 MS1 microscan; MS1 data acquired in profile mode; 15 MS2 dependent scans; 1 MS2 microscan; and MS2 data acquired in centroid mode. MS1 scans acquired in Orbitrap (OT) at 60000 resolution; full MS1 range acquired from 400 to 1500 m/z; MS1 AGC targets set to 1.00E+06; MS1 charge states between 2-5; MS1 repeat counts of 2; MS1 dynamic exclusion durations of 90 sec; ddMS2 acquired in IT; MS2 collision energy and fragmentation: 35% CID; MS2 AGC targets of 1.00E+05; MS2 max injection times of 150 ms.

All samples were analyzed using a 10-step MudPIT sequence.

### MudPIT Data Analysis

Collected MS/MS spectra were searched with the ProLuCID ^46^ algorithm against a database of 73653 protein sequences combining 36636 non-redundant Homo sapiens proteins (NCBI, 2016-06-10 release), 192 common contaminants, and their corresponding 36825 randomized amino acid sequences. In this manuscript, sequences of Halo tag and SNAP tag were added to the database and sequences of AcTEV protease and PreScission Protease were added to contaminants. All cysteines were considered as fully carboxamidomethylated (+57 Da statically added), while methionine oxidation was searched as a differential modification. DTASelect ^47^ v1.9 and swallow, an in-house developed software, were used to filter ProLuCID search results at given FDRs at the spectrum, peptide, and protein levels. Here, all controlled FDRs were less than 1%. Data sets generated from each experiment were contrasted against their merged data set using Contrast v1.9 and in house developed sandmartin v0.0.1. Our in-house developed software, NSAF7 v0.0.1, was used to generate spectral count-based label free quantitation ^48^. For all experiments, only proteins detected in 2 out of total 3 replicates were considered. For single-bait purification MudPIT data, QSPEC ^49^ was used to determine the statistical significance of enriched proteins. Only proteins have FDR<0.05 and Zstatistic value>2 were considered significantly co-purified.

### Imaging Sample Preparation

All cells used for imaging were plated into Mat-Tek dishes with No 1.5 coverslip bottoms. Imaging samples were kept in phenol red free DMEM Medium with GlutaMAX and 10% FBS. While imaging, live cells are kept under 37°C, 5% CO2 and humidified condition. For SPIN1 and SPINDOC imaging, pcDNA5FRT-SNAP-C11orf84 or pcDNA5FRT-SNAP were cotransfected with pcDNA5FRT-Halo-SPIN1 to 293FRT cells. For negative control experiments, pcDNA5FRT-SNAP was co-transfected with pcDNA5FRT-Halo. For positive control experiments, pcDNA5FRT-Halo-NLS-SNAP was transfected. For each experiment, different concentrations of plasmids were transfected to optimize expression level. (data not shown) Cells were imaged 24hr after transfection.

In FRET, HaloTags were stained with HaloTag^®^ Ligands TMRDirect, and SNAP-tags were stained with SNAP-Cell^®^ 505-Star. In SPIN1 and SPINDOC FCCS, HaloTags were stained with HaloTag^®^ Ligands R110Direct, and SNAP tags were stained with SNAP-Cell^®^ 647-SiR. HaloTag^®^ Ligands R110Direct and TMRDirect were added to medium and incubated overnight. Final concentration for R110Direct is 100nM and 50nM for TMRDirect. SNAP-Cell^®^ 505-Star and SNAP-Cell^®^ 647-SiR were added to medium to incubate for 1hr. HOECHST 33258 was also added together with SNAP ligands at a final concentration of 5μg/ml. Cells were then washed for 3 times after SNAP ligands staining with warm culture medium and incubated in fresh medium for at least 30min before imaging.

### Acceptor Photobleaching Förster Resonance Energy Transfer (AP-FRET)

AP-FRET was performed similarly to Weems *et al.* ^50^ In detail, data was acquired with a PerkinElmer Life Sciences UltraVIEW VoX spinning disk microscope controlled by Volocity software. The microscope is equipped with Yokogawa CSU-X1 Spinning disk scanner, both an ORCA-R2 camera (Hamamatsu C10600-10B) and an EMCCD (Hamamatsu C9100-23B) and bleaching studies were conducted with the included PhotoKinesis accessory. The base of the microscope is Carl Zeiss Axiovert 200M. A main dichroic passing reflects 405, 488, 561 and 640nm laser line was used. HaloTag^®^ Ligands TMRDirect labelled proteins were excited by excited by 561nm laser light, and their emission was collected through a dual bandpass 445 (W60), 615 (W70) filter. SNAP-Cell^®^ 505-Star was excited by a 488nm laser and emission was collected through a 525 (W50) bandpass filter. To collect AP-FRET data, time lapse movies were recorded to collect at least 10 timepoints before and after accepter photobleaching. The movies were recorded at a speed of one image per second. For WDR76/SPIN1 experiments, images were recorded using ORCA-R2, using a 40x objective (Oil, NA=1.3). For SPIN1/SPINDOC experiments, images were recorded using EMCCD, and objective was 40x (Water, NA=1.2). For each cell, accepters were bleached by 561nm laser at 100% laser power for 10 cycles. The donor intensity before (*I_Before_*) and after (*I_After_*) were, separately, averaged over time. FRET efficiency (*E*) was represented as: *E* =1-(*I_Before_* / *I_Afer_*). *E* values were calculated in batch using in house imageJ (National Institutes of Health) plugin (accpb FRET analysis jru v1). Control images verified that the acceptor was bleached effectively with the number of iterations.

### Fluorescence Cross-Correlation Spectroscopy (FCCS)

For SPIN1 and SPINDOC FCCS analysis, data was acquired using an LSM-780 (Zeiss) microscope. Cells were imaged with a C-Apochromat 40x (NA=1.2) objective. Green

(HaloTag^®^ Ligands R110Direct) and far red (SNAP-Cell^®^ 647-SiR) fluorophores were employed to eliminate cross-talk between the channels. HaloTag^®^ Ligands R110Direct was excited at 488nm, and its fluorescence was collected through a 491-553nm bandpass filter. SNAP-Cell^®^ 647-SiR was excited at 633nm and its fluorescence was collected through a 633nm longpass filter. For each sample, 10 cells were measured. For both data sets, files were analyzed in Fiji (https://fiji.sc/) using in-house written plugins (analysis cross corr jru v2).

### LC-MS Data Acquisition and Analysis of SCAP-XL Samples

Cross-linked peptides LC-MS3 analysis – Cross-linked peptides were analyzed on an Orbitrap Fusion Lumos mass spectrometer (Thermo Scientific, San Jose, CA) coupled to a Dionex UltiMate 3000 RSCLnano System. Peptides were loaded on the Acclaim™ PepMap™ 100 C18 0.3 mm i.D. 5 mm length trap cartridge (Thermo Scientific, San Jose, CA) with loading pump at 2 μl/min via autosampler. Analytical column with 50 μm i.D. 150 mm length, was packed in-house with ReproSil-Pur C18-AQ 1.9 μm resin (Dr. Masch GmbH, Germany). The organic solvent solutions were water/acetonitrile/formic acid at 95:5:0.1 (v/v/v) for buffer A (pH 2.6), and at 20:80:0.1 (v/v/v) for buffer B. When cross-linked peptides were analyzed, the chromatography gradient was a 20 min column equilibration step in 2% B; a 10 min ramp to reach 10% B; 120 min from 10 to 40 % B; 5 min to reach 95% B; a 14 min wash at 95% B; 1 min to 2% B; followed by a 10 min column re-equilibration step at 2% B. The nano pump flow rate was at 120 nL/min. An MS3 method was made specifically for the analysis of DSSO crosslinked peptides. Full MS scans were performed at 60,000 m/z resolution in the orbitrap with 1.6 m/z isolation window, and the scan range was 375-1500 m/z. Top 3 peptides with charge state 4 to 8 were selected for MS2 fragmentation with 20% CID energy. MS2 scans were detected in orbitrap with 30,000 m/z resolution and dynamic exclusion time is 40 s. Among MS2 fragments, if two peptides with exactly mass difference of 31.9720 with 20 ppm mass tolerance, both of them were selected for MS3 fragmentation at CID energy 35% respectively. MS3 scans were performed in the ion trap at rapid scan with isolation window of 3 m/z, maximum ion injection time was 200 ms. Each MS2 scan was followed by maximum 4 MS3 scans. For the data analysis of DSSO cross-linked peptides Proteome Discoverer 2.2 (Thermo Scientific, San Jose, CA) with add on cross-linking node was used in peptide identification and cross-linked peptide searching. The following settings were used: precursor ion mass tolerance, 10 ppm; fragment ion mass tolerance, 0.6 Da; fixed modification, Cys carbamidomethylation; variable modification, Met oxidation, Lys DSSO Amidated, and Lys DSSO hydrolyzed; maximum equal dynamic modification, 3. Proteins FDR was set at 0.01.

### Cross-linking Data Visualization and Structure Predictions

2D visualization map was generated using xiNET ^30^. Input files for xiNET visualization was directly exported from Proteomics Discoverer. For I-Tasser structure prediction, protein sequences were obtained from Uniprot. Sequences were submitted with or without distance restraints derived from intralinks. For HADDOCK ^37^ docking modeling, SPIN1 structure (4mzf) was submitted with each of the two predicted SPINDOC models and distance restraints derived from interlinks between SPIN1 and SPINDOC. For heterodimer complex docking, the restraints were filtered using DisVis ^51,52^ All structure visualization figures were generated using UCSF Chimera ^53^. All in-house written Fiji or ImageJ plugins can be downloaded at: http://research.stowers.org/imagejplugins/zipped_plugins.html. UCSF Chimera was downloaded from: http://www.rbvi.ucsf.edu/chimera.

### Data availability

The MS dataset may be obtained from the MassIVE database via ftp://massive.ucsd.edu/, MSV000084679, MSV000084713, MSV000084719. Original data underlying this manuscript can be accessed from the Stowers Original Data Repository at LIBPB-1496.

## Supporting information

Movie of Integrative Model of Complex

## Acknowledgements

This body of work is dedicated to the memory of Susan M. Abmayr. Research reported in this publication was supported by the Stowers Institute for Medical Research and the National Institute of General Medical Sciences of the National Institutes of Health under Awards R35GM118068 to JLW and RO1GM112639 to MPW. The content is solely the responsibility of the authors and does not necessarily represent the official views of the National Institutes of Health.

## Author Contributions

X.L., Y.Z., S.M.A, J.L.W, L.F., and M.P.W designed research. X.L., Y.Z., Z.W., Y.H., C.A.S.B., J.L., B.D.S., and J.R.U. performed research. X.L., Y.Z., Z.W., Y.H., C.A.S.B., J.L., B.D.S., J.R.U., and L.F. contributed new reagents/analytic tools. X.L., Y.Z., Z.W., J.L., B.D.S., J.R.U., S.M.A., J.L.W., L.F., and M.P.W. analyzed data. X.L., L.F., and M.P.W wrote the paper.

---

**Figure S1.**
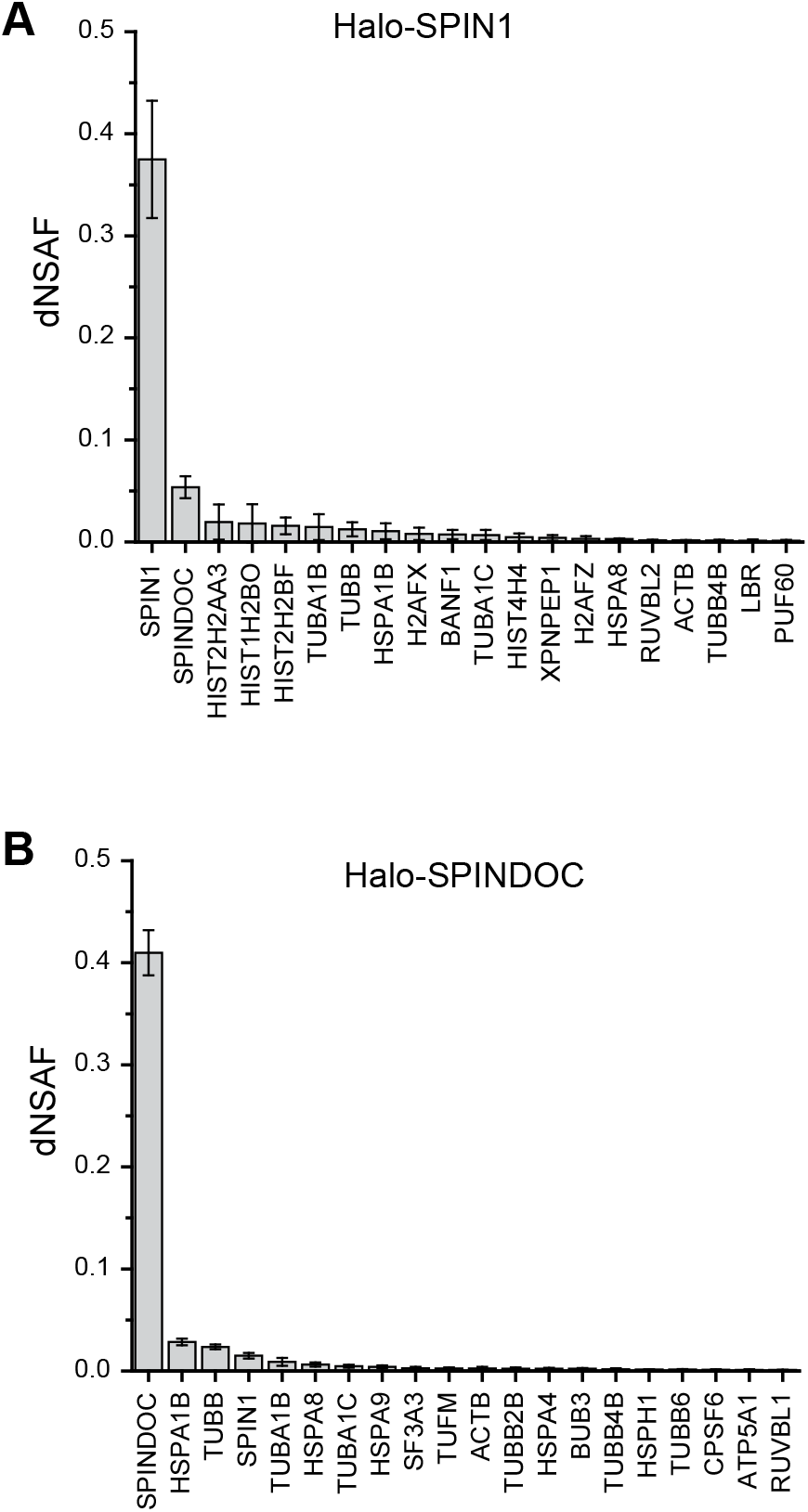
Single Bait Halo-Affinity Purifications. Distributed normalized spectral abundance factor (dSNAF) values with averages and standard deviations measured for three biological replicates are plotted for the top 20 proteins detected in HaloTag-SPIN1 (A) or HaloTagSPINDOC (B) affinity purifications followed by quantitative proteomic analysis (See Supplemental Table S1 for details).

**Figure S2.**
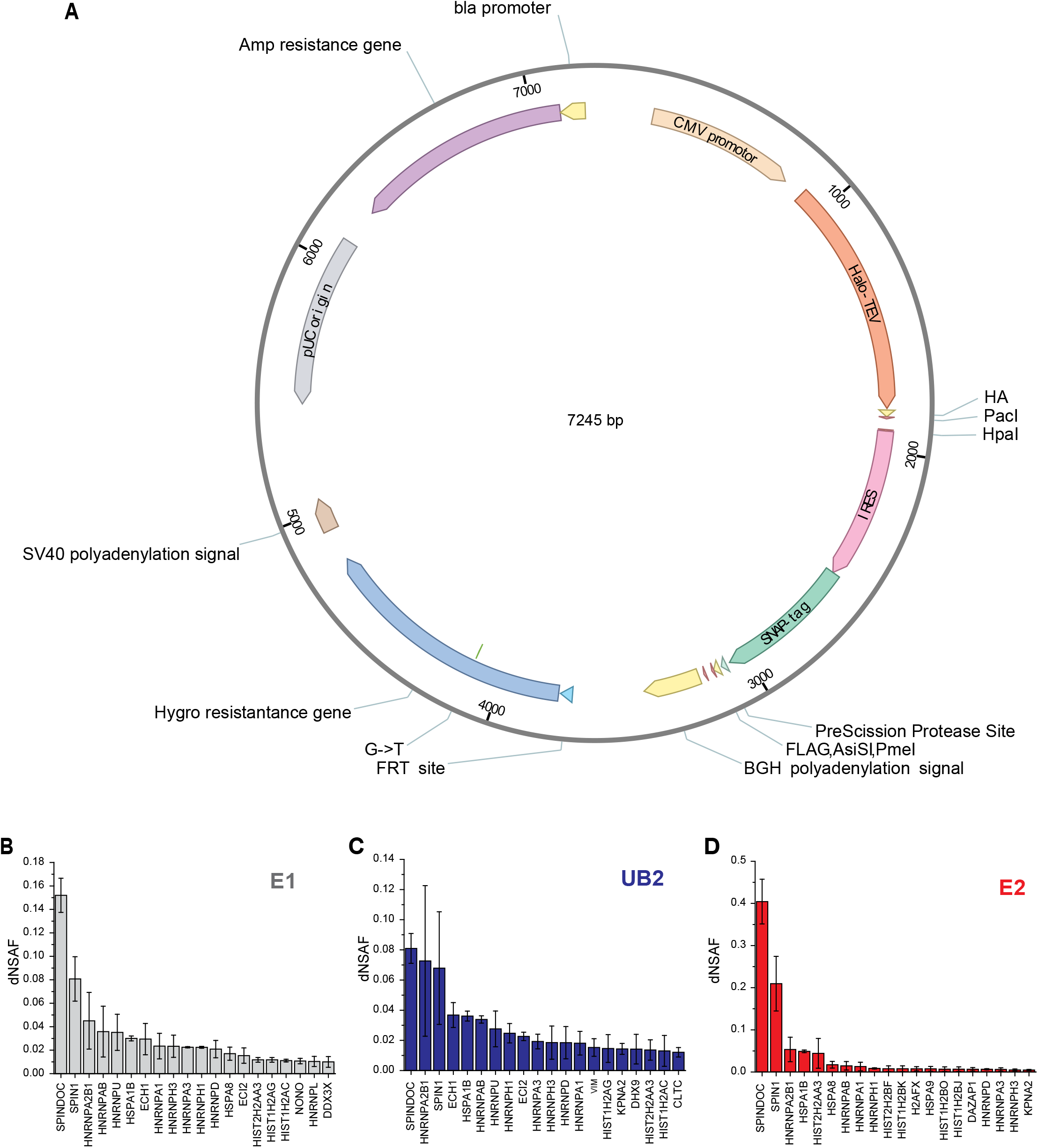
(A) Schematic of pcDNA5/FRT-Halo-TEV-HA-IRES-SNAP-PP-FLAG plasmid used to generate HEK293 double stable cell line. (B-D) The dNSAF plots of the top 20 proteins identified by MudPIT in the E1, UB2, and E2 fractions (See Supplemental Table S2 for details).

**Figure S3.**
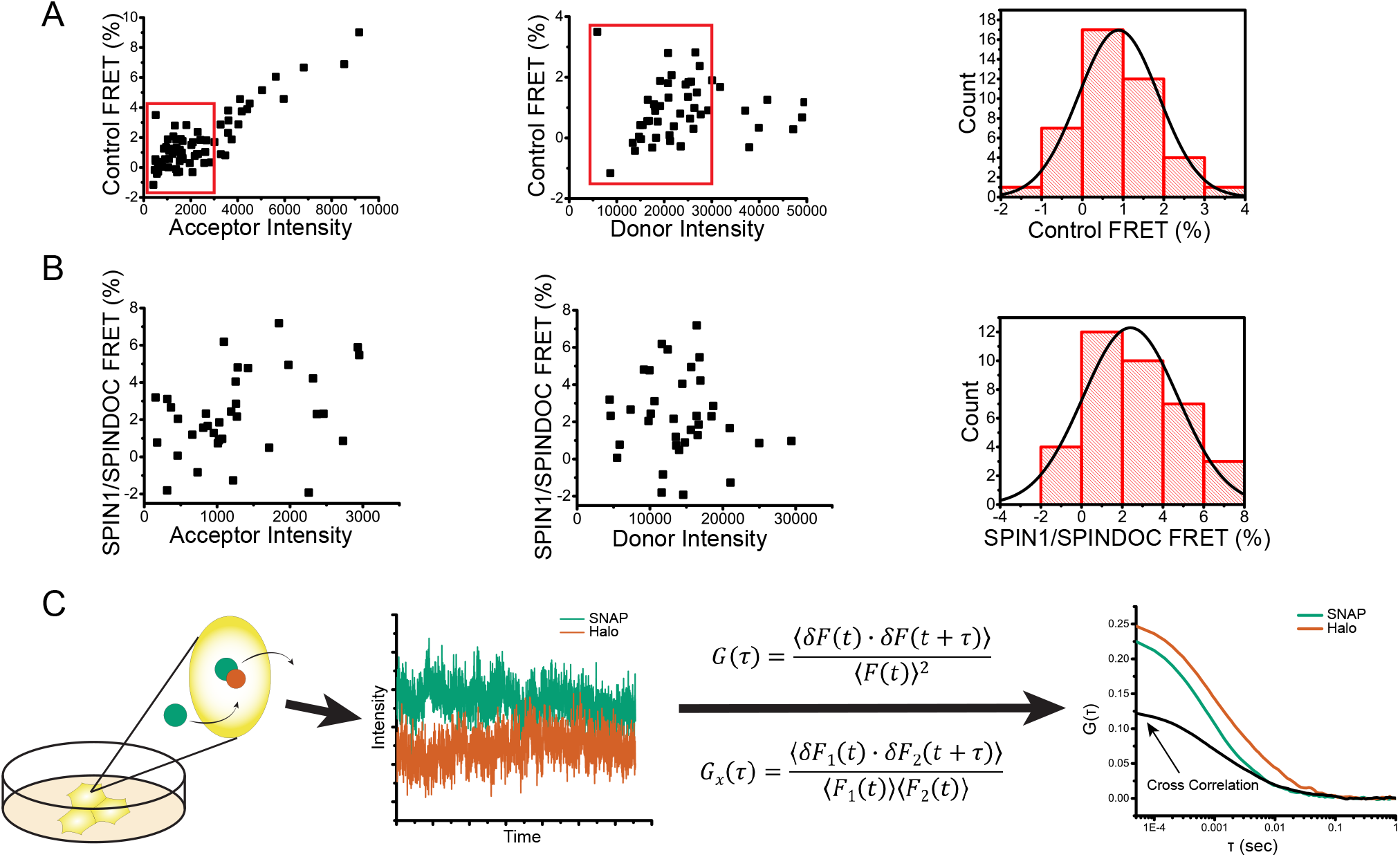
(A) Scatter plot of all Halo-SPIN1 and SNAP-Control FRET data collected (See Supplemental Table S3 for details). Only data points in the red boxes in both donor and acceptor intensity plots were used for the bar charts plotted in Figure 2C. (B) Scatter plot of Halo-SPIN1/SNAP-SPINDOC FRET measurements. The criteria for acceptor and donor intensity cutoff defined in (A) were also applied to Halo-SPIN1/SNAP-SPINDOC FRET measurements to plot the bar graph in Figure 2B. At 0.05 level, data of each sample used for bar chart plotting was tested for normal distribution. (C) Schematic of live cell FCCS method.

**Figure S4.**
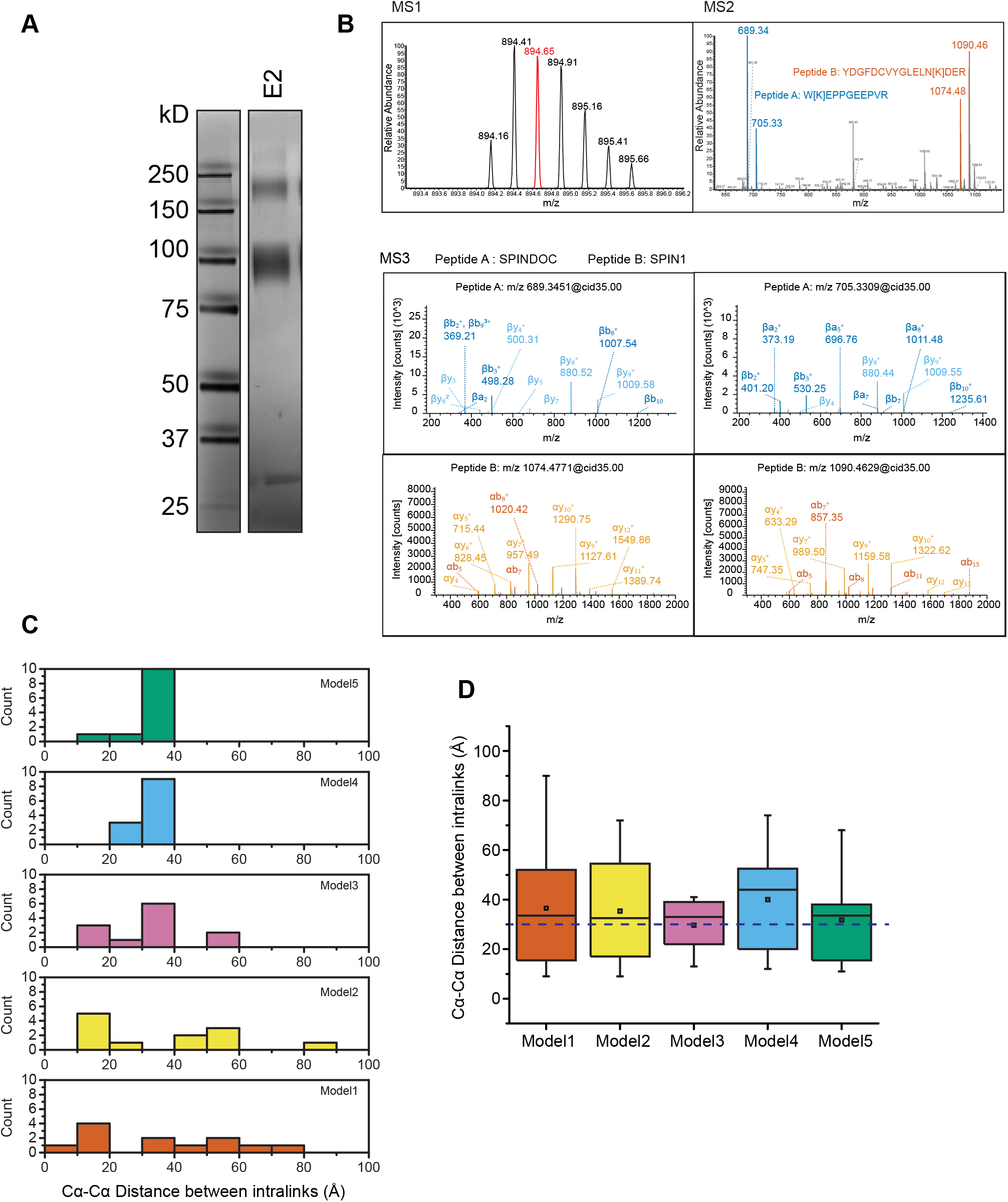
A) Silver-stained SDS-PAGE of SNAP-SPINDOC/Halo-SPIN1 SCAP-XL sample. (B) Example of MS1/MS2/MS3 spectra for two cross-linked peptides. (C) Cα-Cα distances between SPINDOC self-cross-linked sites in structural models predicted by I-Tasser ^34^ (D) Cα-Cα distances between SPINDOC self-cross-linked sites in structural models predicted by I-Tasser ^34^, without inputting any cross-linking information.

## Supplemental Tables

**Table S1:** (A) MS Data Accessibility. (B) Peptide and spectral counts for proteins detected by MudPIT analyses of Halo-SPIN1 (Fig. S1A) and Halo-SPINDOC (Fig. S1B) affinity-purified from HEK293 cells

**Table S2:** Peptide and spectral counts for proteins detected by MudPIT analyses of SCAP fractions E1 (Fig. S2B), UB2 (Fig. S2C), and E2 (Fig. S2D) purified from HEK293 cells

**Table S3:** (A) Raw data for Acceptor Photobleaching Förster Resonance Energy Transfer (AP-FRET) using Halo-SPIN1 as acceptor and SNAP-SPINDOC as donor in live HEK293/FRT cells. (B) Raw values used to calculate self-correlation functions and cross-correlation functions between Halo-SPIN1:SNAP-SPINDOC (Fig. 3D) and Halo-Control:SNAP-SPINDOC (Fig. 3E) **Table S4:** Cross-linked peptides identified from the LC/MS analysis of the SCAP-XL-purified Halo-SPIN1:SNAP-SPINDOC complex (Fig. 4B)

## References

1. Li, T. et al., A scored human protein-protein interaction network to catalyze genomic interpretation. Nature methods 14 (1), 61 (2017).

2. Kotlyar, M., Pastrello, C., Malik, Z., and Jurisica, I., IID 2018 update: context-specific physical protein-protein interactions in human, model organisms and domesticated species. Nucleic acids research 47 (D1), D581 (2019).

3. Sardiu, M. E. et al., Topological scoring of protein interaction networks. Nature communications 10 (1), 1118 (2019).

4. Mellacheruvu, D. et al., The CRAPome: a contaminant repository for affinity purification-mass spectrometry data. Nature methods 10 (8), 730 (2013).

5. Rigaut, G. et al., A generic protein purification method for protein complex characterization and proteome exploration. Nature biotechnology 17 (10), 1030 (1999).

6. Gavin, A. C. et al., Functional organization of the yeast proteome by systematic analysis of protein complexes. Nature 415 (6868), 141 (2002).

7. Los, G. V. and Wood, K., The HaloTag: a novel technology for cell imaging and protein analysis. Methods in molecular biology 356, 195 (2007).

8. Keppler, A. et al., A general method for the covalent labeling of fusion proteins with small molecules in vivo. Nature biotechnology 21 (1), 86 (2003).

9. Bae, N. et al., A transcriptional coregulator, SPIN.DOC, attenuates the coactivator activity of Spindlin1. The Journal of biological chemistry 292 (51), 20808 (2017).

10. Bae, N. et al., Developing Spindlin1 small-molecule inhibitors by using protein microarrays. Nature chemical biology 13 (7), 750 (2017).

11. Chen, W. et al., LINC00473/miR-374a-5p regulates esophageal squamous cell carcinoma via targeting SPIN1 to weaken the effect of radiotherapy. Journal of cellular biochemistry 120 (9), 14562 (2019).

12. Chen, X., Wang, Y. W., and Gao, P., SPIN1, negatively regulated by miR-148/152, enhances Adriamycin resistance via upregulating drug metabolizing enzymes and transporter in breast cancer. Journal of experimental & clinical cancer research : CR 37 (1), 100 (2018).

13. Chew, T. G. et al., A tudor domain protein SPINDLIN1 interacts with the mRNA-binding protein SERBP1 and is involved in mouse oocyte meiotic resumption. PloS one 8 (7), e69764 (2013).

14. Choi, J. W. et al., Spindlin1 alters the metaphase to anaphase transition in meiosis I through regulation of BUB3 expression in porcine oocytes. Journal of cellular physiology 234 (6), 8963 (2019).

15. Devi, M. S. et al., Spindlin docking protein (SPIN.DOC) interaction with SPIN1 (a histone code reader) regulates Wnt signaling. Biochemical and biophysical research communications 511 (3), 498 (2019).

16. Drago-Ferrante, R. et al., Suppressive role exerted by microRNA-29b-1-5p in triple negative breast cancer through SPIN1 regulation. Oncotarget 8 (17), 28939 (2017).

17. Ducroux, A. et al., The Tudor domain protein Spindlin1 is involved in intrinsic antiviral defense against incoming hepatitis B Virus and herpes simplex virus type 1. PLoS pathogens 10 (9), e1004343 (2014).

18. Fagan, V. et al., A Chemical Probe for Tudor Domain Protein Spindlin1 to Investigate Chromatin Function. Journal of medicinal chemistry 62 (20), 9008 (2019).

19. Fang, Z. et al., SPIN1 promotes tumorigenesis by blocking the uL18 (universal large ribosomal subunit protein 18)-MDM2-p53 pathway in human cancer. eLife 7 (2018).

20. Franz, H. et al., The histone code reader SPIN1 controls RET signaling in liposarcoma. Oncotarget 6 (7), 4773 (2015).

21. Gao, Y. et al., Spindlin1, a novel nuclear protein with a role in the transformation of NIH3T3 cells. Biochemical and biophysical research communications 335 (2), 343 (2005).

22. Urh, M., Hartzell, D., Mendez, J., Klaubert, D. H., and Wood, K., Methods for detection of protein-protein and protein-DNA interactions using HaloTag. Methods in molecular biology 421, 191 (2008).

23. Daniels, D. L. et al., Examining the complexity of human RNA polymerase complexes using HaloTag technology coupled to label free quantitative proteomics. Journal of proteome research 11 (2), 564 (2012).

24. Banks, C. A. S. et al., A Structured Workflow for Mapping Human Sin3 Histone Deacetylase Complex Interactions Using Halo-MudPIT Affinity-Purification Mass Spectrometry. Molecular & cellular proteomics : MCP 17 (7), 1432 (2018).

25. Jares-Erijman, E. A. and Jovin, T. M., FRET imaging. Nature biotechnology 21 (11), 1387 (2003).

26. Bacia, K., Kim, S. A., and Schwille, P., Fluorescence cross-correlation spectroscopy in living cells. Nature methods 3 (2), 83 (2006).

27. Kao, A. et al., Development of a novel cross-linking strategy for fast and accurate identification of cross-linked peptides of protein complexes. Molecular & cellular proteomics : MCP 10 (1), M110 002212 (2011).

28. Klykov, O. et al., Efficient and robust proteome-wide approaches for cross-linking mass spectrometry. Nature protocols 13 (12), 2964 (2018).

29. Liu, F., Lossl, P., Scheltema, R., Viner, R., and Heck, A. J. R., Optimized fragmentation schemes and data analysis strategies for proteome-wide cross-link identification. Nature communications 8, 15473 (2017).

30. Combe, C. W., Fischer, L., and Rappsilber, J., xiNET: cross-link network maps with residue resolution. Molecular & cellular proteomics : MCP 14 (4), 1137 (2015).

31. Yang, N. et al., Distinct mode of methylated lysine-4 of histone H3 recognition by tandem tudor-like domains of Spindlin1. Proceedings of the National Academy of Sciences of the United States of America 109 (44), 17954 (2012).

32. Zhao, Q. et al., Structure of human spindlin1. Tandem tudor-like domains for cell cycle regulation. The Journal of biological chemistry 282 (1), 647 (2007).

33. Su, X. et al., Molecular basis underlying histone H3 lysine-arginine methylation pattern readout by Spin/Ssty repeats of Spindlin1. Genes & development 28 (6), 622 (2014).

34. Yang, J. and Zhang, Y., I-TASSER server: new development for protein structure and function predictions. Nucleic acids research 43 (W1), W174 (2015).

35. Zhang, C., Freddolino, P. L., and Zhang, Y., COFACTOR: improved protein function prediction by combining structure, sequence and protein-protein interaction information. Nucleic acids research 45 (W1), W291 (2017).

36. de Vries, S. J., van Dijk, M., and Bonvin, A. M., The HADDOCK web server for data-driven biomolecular docking. Nature protocols 5 (5), 883 (2010).

37. van Zundert, G. C. P. et al., The HADDOCK2.2 Web Server: User-Friendly Integrative Modeling of Biomolecular Complexes. Journal of molecular biology 428 (4), 720 (2016).

38. Wang, W. et al., Nucleolar protein Spindlin1 recognizes H3K4 methylation and stimulates the expression of rRNA genes. EMBO reports 12 (11), 1160 (2011).

39. Wang, J. X. et al., SPINDLIN1 promotes cancer cell proliferation through activation of WNT/TCF-4 signaling. Molecular cancer research: MCR 10 (3), 326 (2012).

40. Chen, X. et al., Suppression of SPIN1-mediated PI3K-Akt pathway by miR-489 increases chemosensitivity in breast cancer. The Journal of pathology 239 (4), 459 (2016).

41. Robaa, D. et al., Identification and Structure-Activity Relationship Studies of SmallMolecule Inhibitors of the Methyllysine Reader Protein Spindlin1. ChemMedChem 11 (20), 2327 (2016).

42. Wagner, T. et al., Identification of a small-molecule ligand of the epigenetic reader protein Spindlin1 via a versatile screening platform. Nucleic acids research 44 (9), e88 (2016).

43. Xiong, Y. et al., Discovery of a Potent and Selective Fragment-like Inhibitor of Methyllysine Reader Protein Spindlin 1 (SPIN1). Journal of medicinal chemistry 62 (20), 8996 (2019).

44. Swanson, S. K., Florens, L., and Washburn, M. P., Generation and analysis of multidimensional protein identification technology datasets. Methods in molecular biology 492, 1 (2009).

45. Florens, L. et al., Analyzing chromatin remodeling complexes using shotgun proteomics and normalized spectral abundance factors. Methods 40 (4), 303 (2006).

46. Xu, T. et al., ProLuCID: An improved SEQUEST-like algorithm with enhanced sensitivity and specificity. Journal of proteomics 129, 16 (2015).

47. Tabb, D. L., McDonald, W. H., and Yates, J. R., 3rd, DTASelect and Contrast: tools for assembling and comparing protein identifications from shotgun proteomics. Journal of proteome research 1 (1), 21 (2002).

48. Zhang, Y., Wen, Z., Washburn, M. P., and Florens, L., Refinements to label free proteome quantitation: how to deal with peptides shared by multiple proteins. Analytical chemistry 82 (6), 2272 (2010).

49. Choi, H., Fermin, D., and Nesvizhskii, A. I., Significance analysis of spectral count data in label-free shotgun proteomics. Molecular & cellular proteomics: MCP 7 (12), 2373 (2008).

50. Weems, J. C. et al., Assembly of the Elongin A Ubiquitin Ligase Is Regulated by Genotoxic and Other Stresses. The Journal of biological chemistry 290 (24), 15030 (2015).

51. van Zundert, G. C. and Bonvin, A. M., DisVis: quantifying and visualizing accessible interaction space of distance-restrained biomolecular complexes. Bioinformatics 31 (19), 3222 (2015).

52. van Zundert, G. C. et al., The DisVis and PowerFit Web Servers: Explorative and Integrative Modeling of Biomolecular Complexes. Journal of molecular biology 429 (3), 399 (2017).

53. Pettersen, E. F. et al., UCSF Chimera--a visualization system for exploratory research and analysis. Journal of computational chemistry 25 (13), 1605 (2004).

